# Localization of Cadherins in the postnatal cochlear epithelium and their relation to space formation

**DOI:** 10.1101/2023.09.30.560287

**Authors:** Holly J. Beaulac, Vidhya Munnamalai

## Abstract

The sensory epithelium of the cochlea, the organ of Corti, has complex cytoarchitecture consisting of mechanosensory hair cells intercalated by epithelial support cells. The support cells provide important trophic and structural support to the hair cells. Thus, the support cells must be stiff yet compliant enough to withstand and modulate vibrations to the hair cells. Once the sensory cells are properly patterned, the support cells undergo significant remodeling from a simple epithelium into a structurally rigid epithelium with fluid-filled spaces in the murine cochlea. Cell adhesion molecules such as cadherins are necessary for sorting and connecting cells in an intact epithelium. To create the fluid-filled spaces, cell adhesion properties of adjoining cell membranes between cells must change to allow the formation of spaces within an epithelium. However, the dynamic localization of cadherins has not been properly analyzed as these spaces are formed. There are three cadherins that are reported to be expressed during the first postnatal week of development when the tunnel of Corti forms in the cochlea. In this study, we characterize the dynamic localization of cadherins that are associated with cytoskeletal remodeling at the contacting membranes of the inner and outer pillar cells flanking the tunnel of Corti.

**Key findings:** F-actin remodeling occurs between E18.5 to P7 in the cochlear sensory epithelium.

Transient changes of F-actin cytoskeleton drives epithelial morphogenesis.

Fluid-filled spaces in epithelium is driven by changes in cell adhesion.

## 1. Introduction

Epithelial morphogenesis requires complex cytoskeletal remodeling and maturation ^1,2^ and is essential for proper organ formation and organ function. Cadherin-driven cell adhesion provides epithelial stability during morphogenesis by inducing cell-to-cell contact-mediated mechanical coupling with the cellular F-actin cytoskeleton ^2,3^. Cadherins are Ca^2+^-dependent cell adhesion molecules that lead to the formation of mature adherens junctions during tissue development ^4^. Initial recruitment of cadherin molecules at the membrane is followed by cadherin clustering and is then stabilized by coupling to the actin cytoskeleton ^5–7^. Maturation of adherens junctions and strengthening of cell-cell adhesion further induces tension-dependent actin remodeling and recruitment of actin binding proteins to the site of de novo actin synthesis^8,9^.

In the cochlea, auditory function is dependent on proper epithelial morphogenesis of the sensory epithelium, the organ of Corti. The organ of Corti attains a highly specialized organization of mechanosensory hair cells intercalated by support cells ^10,11^. At neonatal stages of the mouse cochlea, the organ of Corti has already established its classical mosaic pattern of hair cells intercalated by support cells (Figure 1). There are two classes of mechanosensory hair cells: one row of inner hair cells (IHCs) and three rows outer hair cells (OHCs) (Figure 1A) that are innervated by afferent and efferent neurons enabling the transmission of auditory information along central auditory pathways ^12,13^. The inner and outer hair cells are segregated by the IPC and OPC support cells. The outer hair cells are intercalated by a specific subtype of support cells known as the Deiters’ cells (DCs) (Figure 1A, A’). Thus far, most studies have focused on the cytoskeleton and cytoskeletal remodelers related to stereocilia formation in sensory hair cells; however, the support cells have not received the same attention. Recent studies have shown that the support cells have important electrophysiology, metabolic and sensor properties that are necessary for proper auditory function ^14–16^. The support cells also serve as important support structures for the hair cells ^17^ and help to establish the overall morphology and complex cytoarchitecture of the sensory epithelium. The organ of Corti must be stable enough to withstand vibrational movements of the basilar membrane ^16,18^ and the movement of fluid through the epithelium ^19–21^ to facilitate the propagation of sound information. During sensory epithelial morphogenesis, fluid-filled spaces such as the tunnel of Corti, the Nuel space and the outer tunnel are formed within the epithelium ^22^, which require stabilization of cell and epithelial structure. We show that between late embryonic and early postnatal stages, the support cells undergo significant dynamic morphological changes. However, thus far, no studies have uncovered the spatiotemporal characteristics of cytoskeletal modifications and how the tunnel of Corti is formed during maturation of the cochlear sensory epithelium.

**Figure 1:**
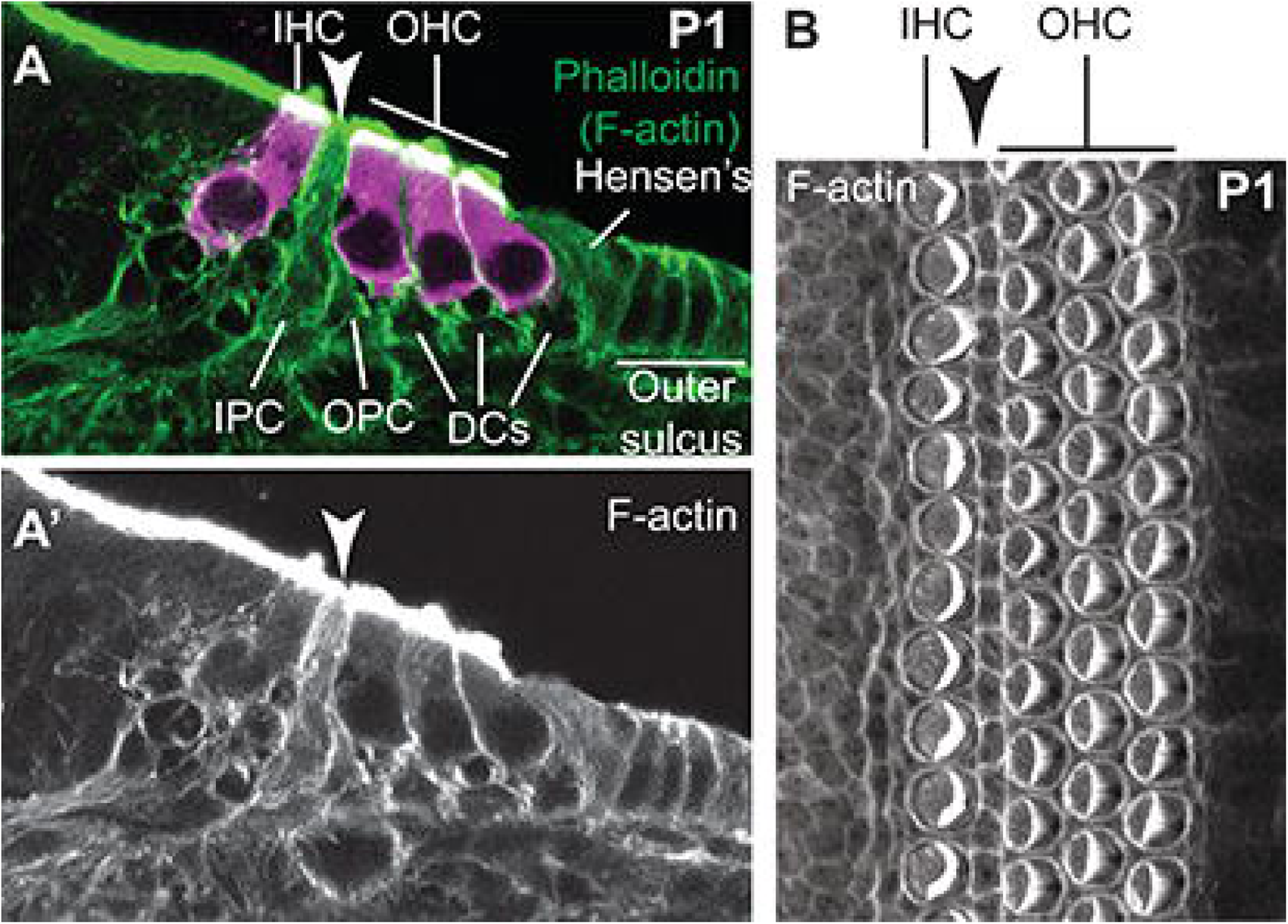
Cross-sectional and top-down view of organ of Corti reveal cytoarchitecture of the support cells and hair cells. (A) Phalloidin labeling of F-actin (green) and Myo6 labeling of HCs (magenta) in a P1 sectioned cochlea shows structural details of the F-actin cytoskeleton in the sensory epithelium along the apico-basolateral axis. (A’) Phalloidin labeling alone. White Arrow indicates the location of the PCs. (B) Top-down view of a Phalloidin-labeled P1 wholemount cochlea shows the overall organization and location of the IHC, PC (black arrow) and the three rows of OHCs.

In this study, we focus on the F-actin cytoskeletal changes that the support cells undergo and the cell adhesion modifications that occur during a specific time in development when fluid-filled spaces form in the sensory epithelium, particularly the pillar cells (PCs) that undergo the most drastic structural cytoskeletal remodeling to form the tunnel of Corti. The tunnel of Corti forms between the IPCs and the OPCs during the neonatal ages of development; however, a close examination of the spatial changes of the F-actin cytoskeleton and cell adhesion of the cochlea undergoing extensive remodeling and the tunnel of Corti opening, has not been performed. Although the F-actin cytoskeleton establishes the structure and shape of all cells ^23^, its expression and structural organization can vary in different cell types in the cochlea. Since epithelial integrity is dependent on cadherins in the cochlea; for example, integrity of the stria vascularis is dependent on cadherins ^24^, we also examine whether the cellular spatial localization of cadherins play a role in establishing the integrity of the sensory epithelium with spaces interspersed within the cochlear sensory epithelium. Since this study focuses on the support cells, the cross-sectional view of the organ of Corti reveals structural details (Figure 1A, A’) that are difficult to appreciate in wholemount cochlear preparations where the stereocilia on hair cells and the overall patterning of the sensory epithelium are the prime focus (Figure 1B). We can also view the immunolabeling in the entire cell along the apico-basolateral axis with finer subcellular differences from the apical, mid- and basolateral regions of the SCs and the membrane versus cytoplasmic localization of proteins. The whole mount provides a single plane view through the epithelium at a time, which may lead to missing out key observations.

Three cadherin molecules reported to be expressed in the mouse cochlea during the first postnatal week are neural (N-), epithelial (E-), and placental (P-) cadherins ^25–27^. The tunnel of Corti forms between the inner pillar cells (IPC) and the outer pillar cells (OPCs), spaces of Nuel and the outer tunnel ^27^. A recent study suggested that the abnormal premature enlargement of the tunnel of Corti was caused by loss of P-cadherin. From the study’s wholemount images, P-cadherin localization in the PCs appeared to be apical ^25^ but was not clear from single optical planes. Cross-sectional views of cadherin localization along the apico-basolateral axis during the narrow temporal window as the tunnel of Corti forms would provide a clearer view of how E-cadherin or N- or P-cadherin localization influence tunnel formation under normal developmental conditions. In this study, we analyze the spatiotemporal localization of these three cadherins in the cochlea.

## 2. Results and Discussion

To identify the correct stage to observe the opening of the tunnel of Corti and characterize the associated subcellular changes in the F-actin cytoskeletal organization, we first analyzed sectioned cochlear tissue on E18.5, P6 and P8. On E18.5, Phalloidin labeling shows that F-actin is not distinguishable in the pillar cells (dotted lines) but is clearly present at the membranes adjoining both the OHCs and the DCs (Figure 2A). Thus, we examined markers for cell adhesion along the OHC-DC membrane contacts. A prior study showed an increase in junctional E-cadherin in the apical junctions of the PCs in wholemount mouse cochleas during the first postnatal week after birth ^28^. Thus, E-cadherin signaling is a strong candidate for PC remodeling. Comparing F-actin with E-cadherin, we found E-cadherin localized to the OHC-DC membrane contacts (Figure 2A’). On E18.5, E-cadherin was also localized to the membranes of Hensen’s cells (Figure 2A’, green arrow). At this stage, there was no specific enrichment of F-actin or E-cadherin in the PCs. Cell adhesion induces bi-directional signaling that promotes the recruitment of phosphorylated actin binding proteins at their tyrosine residues (pTyr), a sign of active actin assembly ^29,30^. Immunolabeling for pTyr was undetectable on E18.5, presumably because minimal remodeling of the F-actin cytoskeleton was required at this stage of development (Figure 2A”). There was no notable patterns of phalloidin, E-cadherin, or pTyr labeling in the PCs on E18.5 (Figure 2A”’).

**Figure 2:**
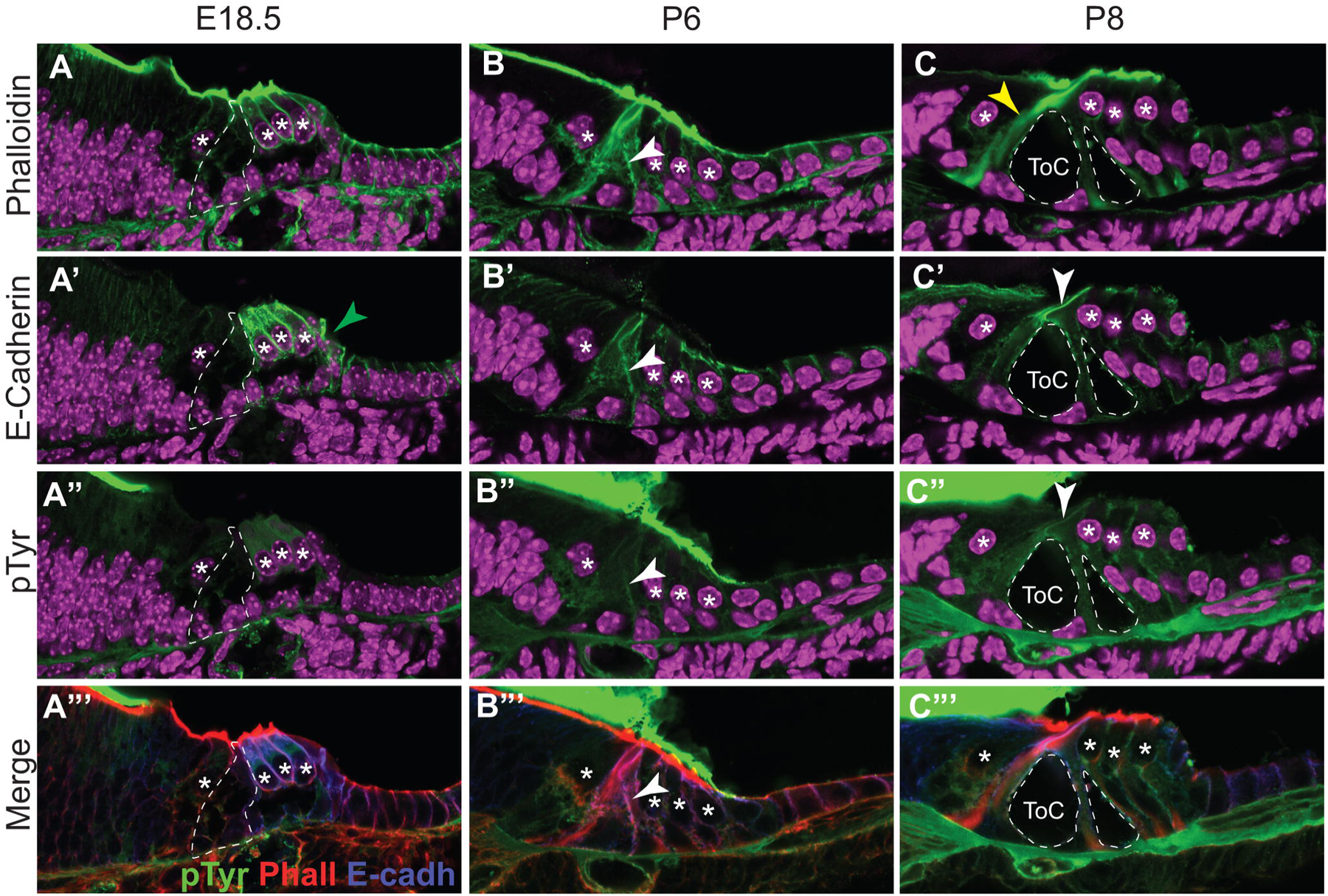
Significant remodeling of F-actin and E-cadherin occurs in the cochlear sensory epithelium between late embryonic and early postnatal ages. (A-A”’) Phalloidin labeling of F-actin, and E-cadherin and pTyr immunolabeled proteins and merged image in E18.5 cochleas. Dotted lines outline the PCs, asterisks denote the HCs. (B-B”’) Phalloidin labeling of F-actin, and E-cadherin and pTyr immunolabeled proteins and merged image in P6 cochleas. Arrows indicate the mid-regions of IPC-OPC membranes that will form the axial regions of the mature PCs. (C-C”’) Phalloidin labeling of F-actin, and E-cadherin and pTyr immunolabeled proteins and merged image in P6 cochleas. Yellow arrow in C indicates actin bundle in IPC. White arrows in C’ and C” indicate the apical regions of IPC-OPC membrane contacts. Dotted lines outline the tunnel of Corti and the space of Nuel. Nuclei are counter stained with DAPI (magenta).

Next we analyzed cochleas on P6. The tunnel of Corti was still not formed at this stage. However, actin filaments were now visible in the PCs, especially in the IPC and the IPC-OPC membrane contacts (Figure 2B, arrow indicates mid-region of IPC-OPC membrane). There is a 5e2 ± 0.95e2 % enrichment of F-actin at the membrane contacts of the mid-IPC-OPC region relative to the cytoplasm (T-test, p = 1.2e-6). E-cadherin was also localized at the adjoining IPC-OPC membranes, but E-cadherin was expressed from the apical to the mid-regions (arrow) along the apico-basolateral axis of the IPC-OPC membranes, indicating there was an enrichment in cell adhesion contacts associated correlating with F-actin in the mid-region of the PCs (Figure 2B’, arrow). We detected an enrichment of 2.9e2 ± 0.52e2 % in the mid-region of IPC-OPC membrane contacts relative to the cytoplasm, assayed at the same region of interest as phalloidin (p = 3.1e-6). At P6, there was minimal recruitment of pTyr actin binding proteins at the IPC-OPC membrane contacts by 1.4e1 ± 0.31e2 percent relative to the cytoplasm, assayed at the same region of interest as phalloidin p = 0.0004). This suggests that there may be a temporal delay in pTyr protein recruitment once E-cadherin localization stabilizes (Figure 2B”). Merged image show co-localization of Phalloidin with E-cadherin (Figure 2B”’).

We then performed the same analysis in the P8 cochlea. By P8, the tunnel of Corti has opened (Figure 2C). There is a clear F-actin bundle that has formed within the IPC (Figure 2C, yellow arrow). However, by P8, F-actin and E-cadherin were no longer identifiable in the mid-regions of the IPC-OPC contact membranes that were now occupied by the tunnel of Corti and Nuel space. Despite the formation of the tunnel of Corti, the IPC and the OPC remain connected at their adjoining apical junctions where F-actin and E-cadherin remains localized (Figure 2C, 2C’, arrow). P-Tyr labeling was not present in the PCs at P8, suggesting that a critical window was missed (Figure 2C”, 2C”’). The time interval between P6 and P8 was too large to observe the transient cytoskeletal modifications associated with the formation of the tunnel of Corti.

To capture the opening of the tunnel of Corti, we harvested P7 cochleas at 3 timepoints throughout the day in the morning (9 am), mid-day (1 pm) and in the evening (5 pm) (Figure 3). In the morning, the tunnel of Corti (ToC) was not yet opened (closed). We observed an accumulation of F-actin at the mid-region of the IPC-OPC membranes (Figure 3A, arrow) along the apico-basolateral axis, which will go onto to become the axial regions of the PCs that flank the tunnel of Corti. We detected an enrichment of phalloidin labeling by 26.7e2 ± 8.9e2 % at the mid-region of IPC-OPC membrane contacts relative to the cytoplasm (p= 5.9e-9). Coincidently, in the same region of interest where we detected phalloidin enrichment, we also saw increases in E-cadherin (Figure 3A’, arrow) and pTyr labeling (Figure 3A”, arrow) at the IPC-OPC membrane contacts with enrichments of 23.6e2 ± 12.2e2 % (p = 5.9e-5) and 13.7e2 ± 3.8e2 % (p = 4.1e-6) respectively relative to the cytoplasm. This indicates that F-actin, E-cadherin and pTyr were co-localized at the same mid-region of interest at the IPC-OPC membranes (Figure 3A”’ inset, arrow). These enrichments of phalloidin, E-cadherin and pTyr at the IPC-OPC membranes relative to the cytoplasm were higher in the closed tunnel on P7 than on P6 when the tunnel was also closed, and E-cadherin and pTyr localization at the membrane were only 2.9 -fold and 1.4 – fold higher than in the cytoplasm respectively (Figure 2B’-B”’). There was some pTyr labeling at the apical junction that the mid-region (Figure 3A”), which suggests that there may be some active F-actin assembly on the apical junctions of the PCs. Unlike the mid-apical region of the IPC-OPC membrane contacts, there was no overlay or significant enrichment of F-actin, E-cadherin and pTyr signals in the basolateral end of the PC membranes relative to cytoplasmic levels (Figure 3A-A”’). We also observed pTyr labeling when the Nuel space has a very narrow opening, which suggests there was also recruitment of actin binding proteins during initiation of Nuel space formation (Figure 3A”, red arrow). At the mid-timepoint on P7, the tunnel of Corti was open at less than 100 μm^2^ in cross-sectional area (small ToC). F-actin and E-cadherin expression was not detectable in the mid-axial region of the IPC-OPC membranes but are present at the apical junction (Figure 3B-B’). Interestingly, pTyr labeling remained high at the edges of cell membranes (Figure 3B”-B”’). This suggests that these sites remained enriched for actin binding proteins which help remodel cellular structure of the tunnel. In the final timepoint on P7, the tunnel was large at more than 120 μm^2^ (large ToC). F-actin and E-cadherin remained at the apical regions of the PCs (Figure 3C’C’); while pTyr labeling was relatively minimal at the apical junctions (Figure 3C”-C”’). This suggests that there was a greater recruitment of pTyr-labeled actin binding proteins where the cochlear spaces are forming and not at stable contacts. The variable localization of E-cadherin on the adjoining IPC-OPC membranes along the apico-basolateral axis suggests that a reduction in cell adhesion precedes the opening of the tunnel of Corti at the mid-region of the PC membranes and was accompanied by a large recruitment of pTyr-labeled proteins. This data also show that P7 is critical timepoint when the support cells in the sensory epithelium undergo significant cellular remodeling to attain its complex cytoarchitecture. Conversely, stabilized F-actin will require less recruitment of actin binding proteins as there are no sites for de novo F-actin synthesis e.g., in the apical junctions of the PCs.

**Figure 3:**
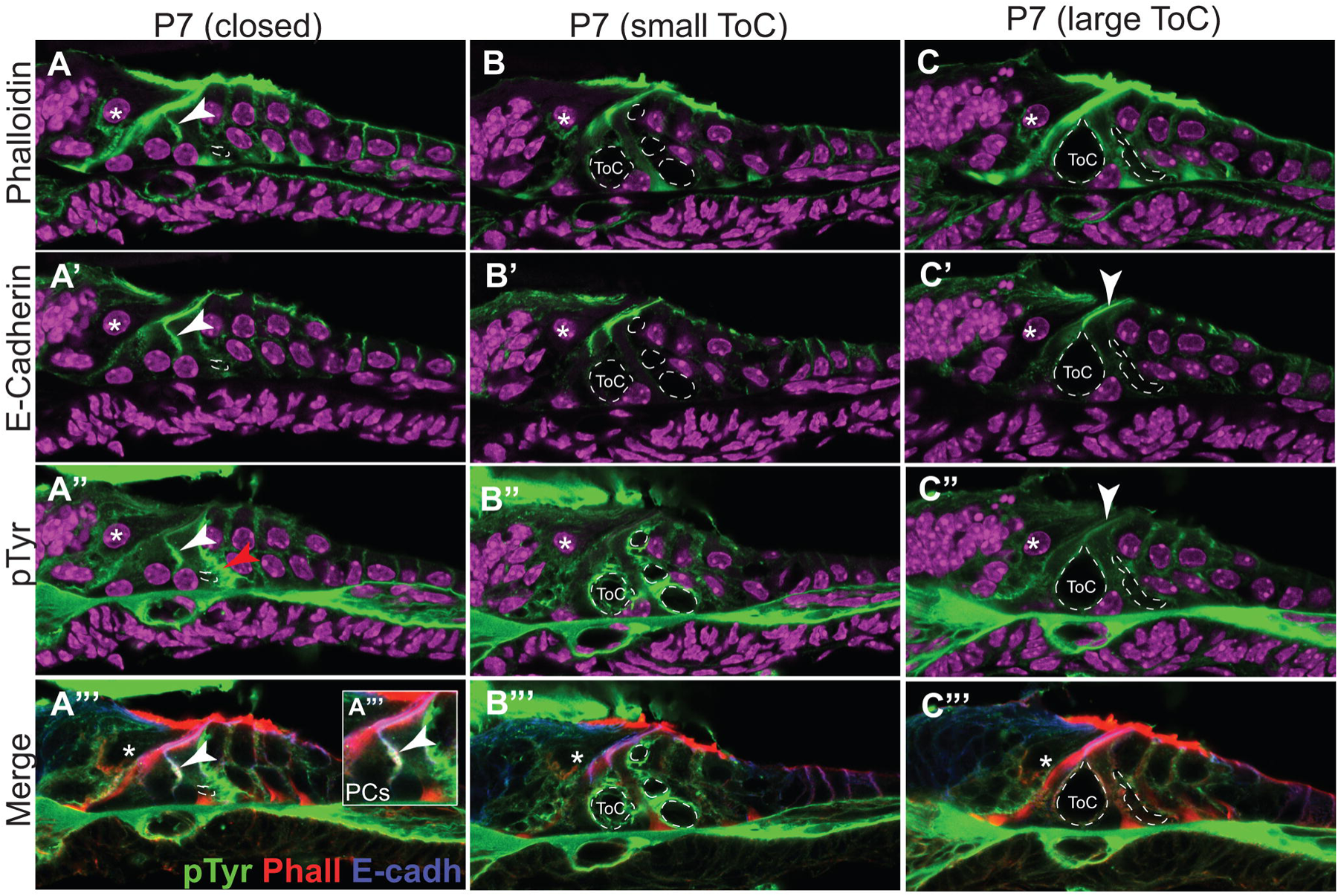
Fine temporal resolution of the opening of the tunnel of Corti reveals novel cytoskeletal details. (A-A”’) Phalloidin labeling of F-actin, and E-cadherin and pTyr immunolabeled proteins and merged image in P7 cochleas harvested at 9 am. Arrows indicate the Arrows indicate the mid-regions of IPC-OPC membrane contacts. A”’ Inset-overlay of mid-region of IPC-OPC junction. E-cadherin (green), pTyr (red) and Phalloidin (blue). White arrow indicates co-localization of the three markers at the mid-region of the IPC-OPC junction. Red arrow indicates pTyr accumulation at OPC-DC contact flanking Nuel space. Dotted lines denote the locations of spaces that will form the tunnel of Corti and the space of Nuel (B-B”’) Phalloidin labeling of F-actin, and E-cadherin and pTyr immunolabeled proteins in P7 cochleas harvested at 1 pm. Dotted lines denote the locations of spaces that will form the tunnel of Corti and the space of Nuel. (C-C”’) Phalloidin labeling of F-actin, and E-cadherin and pTyr immunolabeled proteins and merged image in P7 cochleas harvested at 5 pm. Arrows indicate the apical junctional contacts between the IPCs and the OPCs. Asterisks denote the IHCs, and nuclei are counter stained with DAPI (magenta).

N-cadherin is another cadherin known to be expressed in the developing cochlea ^26^. We examined whether N-cadherin is spatially positioned to influence the opening of the spaces in the cochlear sensory epithelium. On P6, N-cadherin is visible in the inner hair cell (IHC), inner phalangeal and inner border cell, but is clearly absent in the PCs and the DCs (Figure 4A-A’). This expression pattern remains consistent in P7 cochleas at the same three timepoints when the tunnel of Corti was still closed (Figure 4B-B’), has a small opening (Figure 4C-C’) and when the tunnel opening was larger (Figure 4D-D’). Thus, the stable expression of N-cadherin and its consistent absence of expression in the PCs suggests that N-cadherins do not influence the opening of the tunnel of Corti.

**Figure 4:**
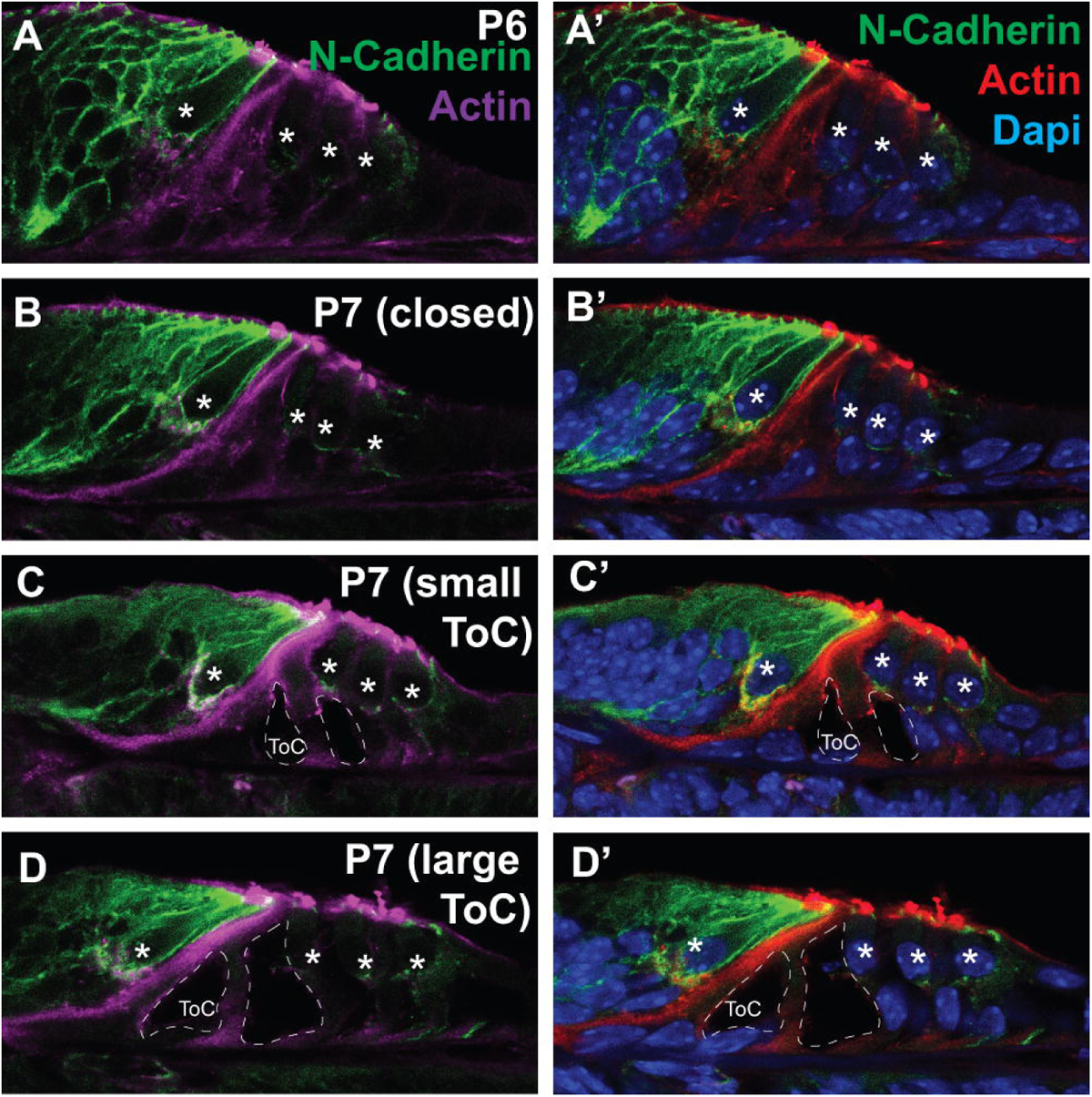
N-cadherin localization remains stable during postnatal ages. (A-A’) On P6, N-cadherin is localized to the membranes of the inner border cell, inner phalangeal cell and the IHC. Merged image is counterstained with DAPI. (B-D) On P7, N-cadherin is localized to membranes of the inner border, IHC and inner phalangeal cell membranes remains stable as the tunnel of Corti is formed. Asterisks denote the hair cells and actin is immunolabeled in magenta. Merged image is counterstained with DAPI.

A recent study showed that P-cadherin is expressed in the PCs of the cochlea during early postnatal stages. Upon Thyroxine administration, P-cadherin was translocated from the PC membranes to the cytoplasm, thereby reducing cell-cell adhesion ^25^. The study analyzed wholemount cochleas, showing images from a single plane; thus, the expression of P-cadherin along the entire apico-basolateral axis of PCs was unclear. To determine its spatial localization, we analyzed P-cadherin expression. On P6, P-cadherin was localized in the IPCs, but enriched at the top apical end of the IPC (Figure 5A-A’, white arrow). In addition to the PCs, we found P-cadherin was also expressed in the membranes of the Hensen’s cells and the cells in the outer sulcus region (Figure 5A, green arrow). The expression of P-cadherin was much more substantial in these lateral non-sensory cells compared with E-cadherin. On P7, while the tunnel of Corti was still closed, P-cadherin remained enriched in the apical portion of the PCs, the Hensen’s cells and the outer sulcus cells (Figure 5B-B’). The apical spatial localization remained unchanged while the tunnel of Corti opened (Figure 5C, 5C’, 5D, 5D’). The expression of P-cadherin remained stable in the Hensen’s and outer sulcus cells as well.

**Figure 5:**
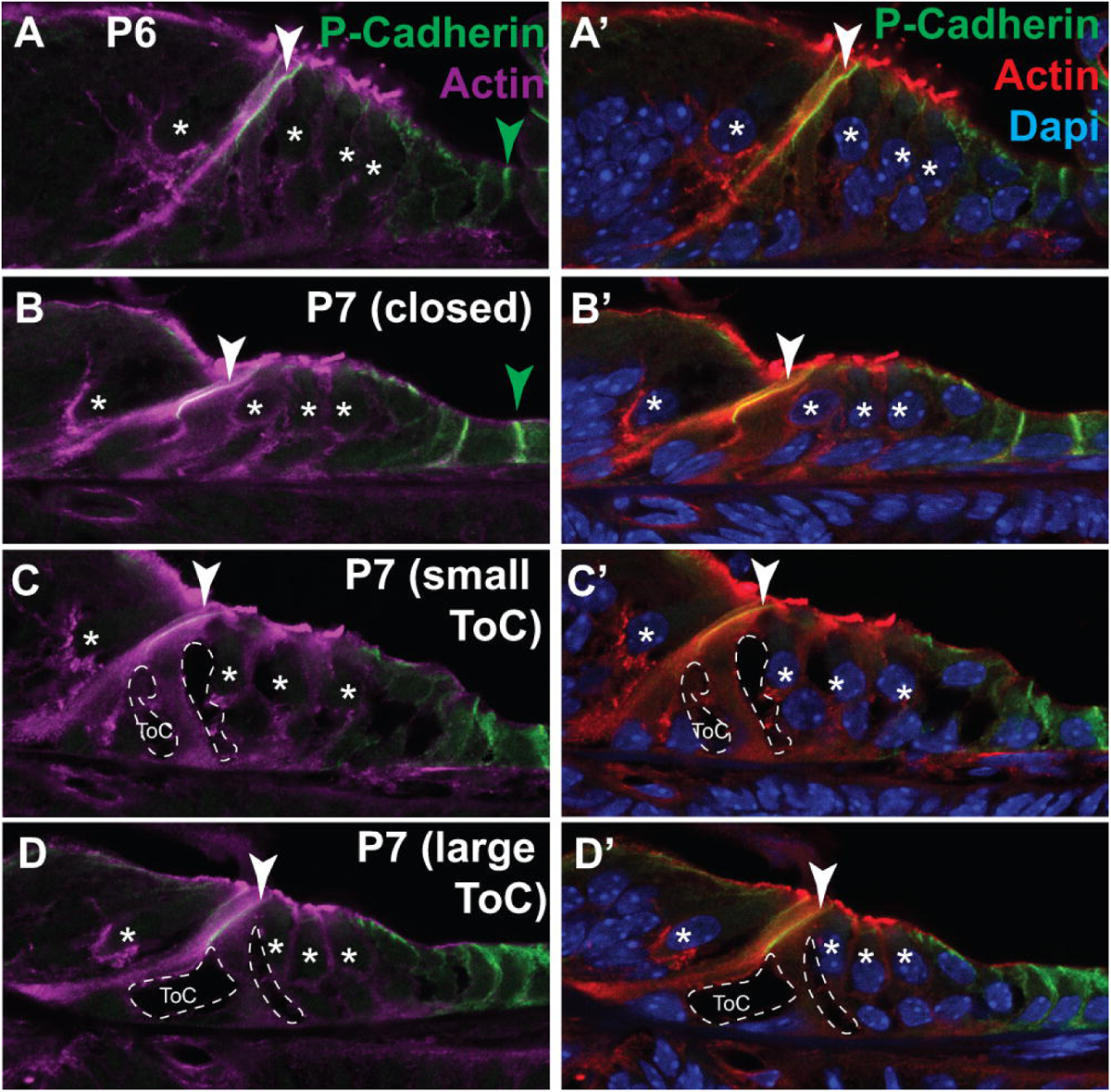
P-cadherin localization remains stable during postnatal ages. (A-A’) On P6, P-cadherin is localized to the membranes of the IPC (white arrow), and Hensen’s and outer sulcus cells (green arrow). Merged image is counterstained with DAPI. (B-D) On P7, P-cadherin is localized to the membranes of the IPCs and the OPCs, Hensen’s cells and outer sulcus cells. Arrows indicate the apical portion of the IPC-OPC adherens junctions. Asterisks denote the hair cells, and actin is immunolabeled in magenta. Merged image is counterstained with DAPI.

Since pTyr labeled proteins were recruited to sites of F-actin and E-cadherin accumulation at IPC-OPC membrane prior to the opening of the tunnel of Corti (Figure 3). We postulated that in preparation for the tunnel opening, actin binding proteins are recruited to facilitate the remodeling of F-actin in the PCs. Thus, we examined the expression of Cofilin, an F-actin severing protein, on P6 and the three stages of tunnel of Corti formation on P7 (Figure 6). Previous studies showed cofilin labeling in the HCs ^31,32^; however, Cofilin localization in the SCs has not been characterized thus far. Thus, we were surprised to find punctate labeling of Cofilin in the cytoplasm of the SCs but not in the HCs (Figure 6A-D). On P6, we saw a 70 ± 10 % (p = 5.5e-7) enrichment of Cofilin localized in the mid-region of the IPC-OPC junction relative to cofilin expression in the cytoplasm of the PCs (Figure 6A-A”). On P7, prior to opening of the tunnel, we saw a 320 ± 40 % (p = 6.2e-7) enrichment of Cofilin localized in the mid-region of the IPC-OPC junction relative to cofilin expression in the cytoplasm of the PCs (Figure 6B-B”, arrow). This suggests that during this critical period, Cofilin recruitment is associated with the actin remodeling and aids in disrupting cell adhesion in the mid-region of the IPC-OPC membranes.

**Figure 6:**
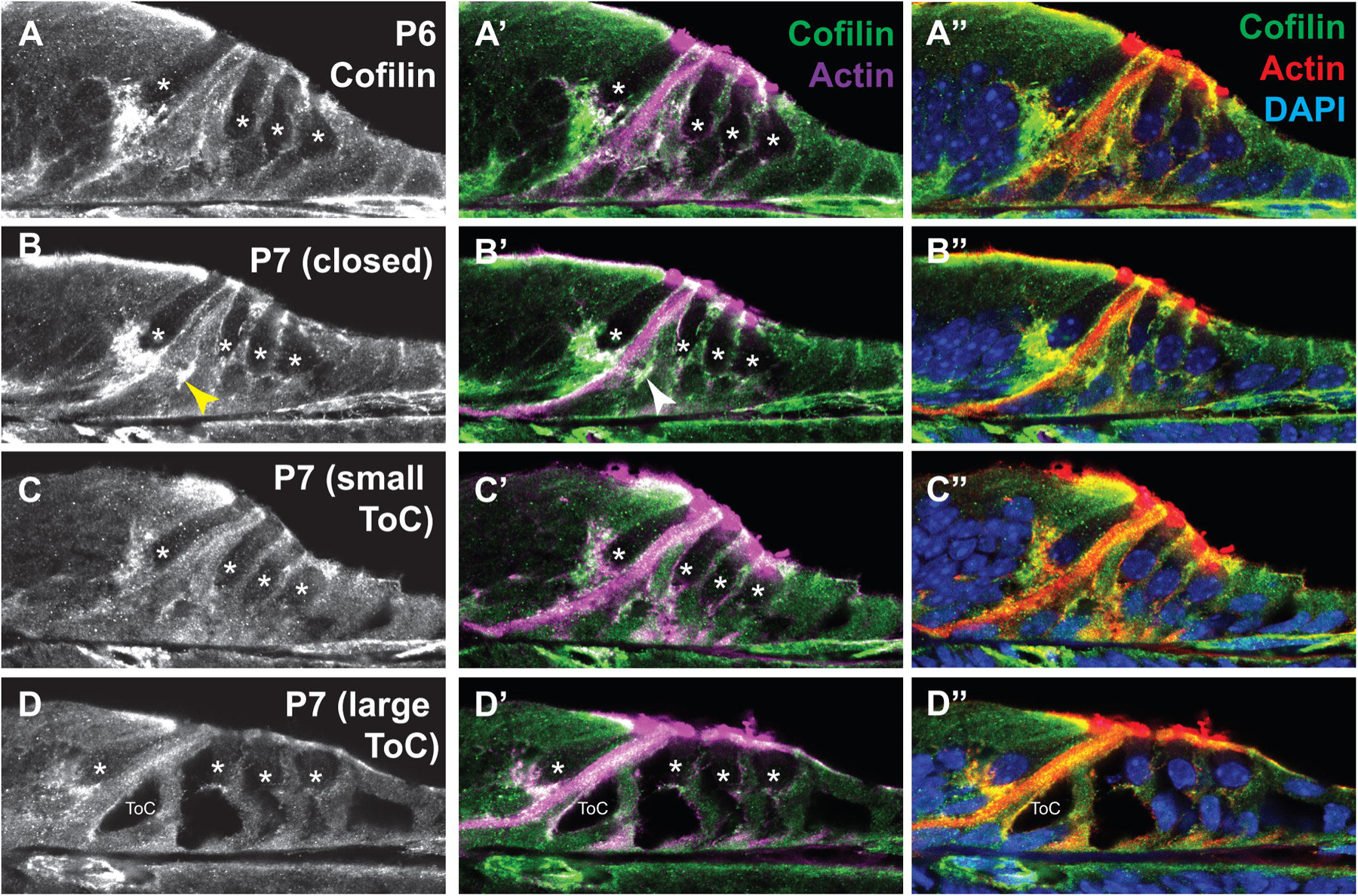
Cofilin is expressed in the support cells of the sensory epithelium in the postnatal cochlea. (A-A”) On P6, Cofilin labeling is punctate in the cytoplasm and high at the PC membranes. Cofilin is expressed in the SCs, but not the HCs. Merged image is counterstained with DAPI. (B-B”) On P7, before the tunnel of Corti forms, Cofilin is present in the mid-region of the IPC-OPC junctions (arrow). Merged image is counterstained with DAPI. (C-D) As the tunnel opens, Cofilin remains at the edge of the cell membrane that retracts to form the tunnel of Corti. Merged image is counterstained with DAPI. Asterisks denote the hair cells, and actin in immunolabeled in magenta. Yellow arrow indicates cofilin enrichment.

The stable P-cadherin expression in the apical portion of the PCs is unlikely to influence the opening of the tunnel of Corti under normal developmental conditions. On the other hand, the dynamic and transient localization of E-cadherin in the mid-regions of the IPC-OPC membranes, and its subsequent downregulation preceding the opening of the tunnel of Corti makes the dynamic downregulation E-cadherin a much more likely mechanism for opening the tunnel under normal developmental conditions. However, it is possible that in a disease context, the loss of P-cadherin can exacerbate the tunnel opening and final size of the tunnel, disrupting the overall stability of the sensory epithelium. Perhaps both E-cadherin and P-cadherin trafficking are impacted when the cytoarchitecture of the sensory epithelium is disrupted under disease conditions. Thus, follow up studies are needed to validate this. In this study, we show evidence that significant actin remodeling takes place in the SCs during P6-P8 and paves way for future investigations on actin assembly in the SCs on which little focus has been placed thus far. Here we closely analyze the cell adhesion characteristics associated with the opening of the tunnel of Corti. We predict that the same decrease in cadherin trafficking will promote the formation of the Nuel space and the outer tunnel. Catenins are intracellular cell adhesion proteins that link cadherins to the F-actin cytoskeleton ^33^; thus, are critical players in stabilizing adherens junctions and assembling epithelia, including the cochlear epithelium ^26,28,34^. However, how their localizations are modified during the critical stages of the opening of cochlear spaces would be a good focus for future studies.

## 3. Experimental Procedures

### 3.1 Mice

Time-mated Swiss Webster dams (Envigo, Indianapolis, IN, USA) were denoted as embryonic day (E)0.5 on the first day a plug was observed. E18.5 embryos, and postnatal day (P)6-P8 pups were euthanized by decapitation, in accordance with the Institutional Animal Care and Use Committee (IACUC) guidelines at The Jackson Laboratory, Bar Harbor, and heads or ears were harvested and fixed in 4% paraformaldehyde (PFA) (Electron Microscopy Sciences, Hatfield, PA, USA).

### 3.2 Histology

After fixation, E18.5 heads were bisected and processed through a 10%, 20%, and 30% sucrose gradient series, embedded with TFM and cryofrozen. For postnatal studies, cochleas were extracted on P6, P7 and P8, and fixed in 4% PFA for three hours at 4°C, immersed in a 30% sucrose and transferred to a 1:1 solution of 30% sucrose: Tissue Freezing Medium (TFM) (General Data Healthcare, Cincinnati, OH, USA) before final embedding in 100% TFM and cryofrozen. Tissues were cryosectioned at 12 μm for histology at all ages. To capture the opening of the tunnel of Corti (ToC) on P7, inner ears were harvested and fixed at 3 timepoints throughout the day in the morning (9 am), mid-day (1 pm) and in the evening (5 pm).

### 3.3 Immunofluorescence

Tissues were washed in PBS/0.5% Triton and blocked at for at room temperature (RT) 1 hour with 2% Donkey serum (Jackson ImmunoResearch, West Grove, PA, USA) in PBS/0.5% Triton, followed by overnight primary antibody incubation at 4°C. Primary antibodies included: rat anti-E-cadherin (1:250, AB11512, Abcam, Waltham, MA, USA), sheep anti-N-cadherin (1:250, AF6426, R&D Systems, Minneapolis, MN, USA), goat anti-P-cadherin (1:250, AF761, R&D Systems), mouse anti-phosphoTyrosine (1:250, zms1b282, Sigma-Aldrich), anti-Cofilin (1:250, 5175, Cell Signaling Technology, Danvers, MA, USA). For N-cadherin, P-cadherin and Cofilin immunolabeling, antigen retrieval was performed prior to primary antibody incubation with a 10 mM sodium citrate/ 0.05% Tween20 buffer at 6.0 pH for 15 min at 99°C. Since phalloidin labeling of F-actin is incompatible with antigen retrieval, we co-labeled with an actin antibody (1:250, MA5-11869, Invitrogen). Alexa-conjugated secondary antibodies (Invitrogen) were incubated for 2 hours at RT. Nuclei were counter stained with DAPI (Abcam, Waltham, MA, USA) and tissues were mounted with Fluoromount G mounting medium (Life Technologies Corporation, Carlsbad, CA, USA).

### 3.4 Data Acquisition, Analysis and Image processing

60X magnification images were acquired on a Zeiss LSM 800 confocal microscope (ZEISS, Oberkochen, BW, Germany) in The Jackson Laboratory microscopy core. Raw data were analyzed using the FIJI software (NIH). At the morning time point, mid-turns were consistently closed. In themed-day timepoint, we qualified small ToCs as those tunnels with a range of cross-sectional areas between 35 μm^2^ – 100 μm^2^. In the afternoon timepoint, ToCs were consistently open, and we qualified large ToCs to be greater than 120 μm^2^. Data were collected and quantified on 2 sections per cochlea for n = 6 cochleas at each time point. All 6 cochleas in each of the presented figures were immunolabeled and data were acquired at the same time and settings. To quantify enrichment of signals, a 0.6 μm^2^ region of interest was used to measure levels at the IPC-OPC membrane and the IPC cytoplasm within the same sample. We utilized the same region of interest at the same coordinates across channels to define co-localized signals.

We calculated the percent difference between the membrane versus the cytoplasm. To qualify specific localization at the membrane, student’s t-test were performed to determine significant differences in levels at the membrane versus the cytoplasm. Bon-Ferroni correction factor was applied for multiple comparisons. TIFF images for figures were processed in Adobe Photoshop and assembled in Adobe Illustrator (Adobe, San Jose, CA, USA).

## Acknowledgements

The authors acknowledge the Microscopy Core at The Jackson Laboratory for equipment maintenance and upkeep, and assistance. The authors thank members of the Munnamalai lab for providing critical feedback on the manuscript.

## 5. Author Contributions

H.J.B., study conceptualization and design, development of methodology, acquisition, data analysis, writing of the manuscript. V.M., funding, material support, study conceptualization and design, development of methodology, acquisition, analysis, and interpretation of data; writing and revision of the manuscript. All authors read and approved the final manuscript.

## 6. Ethics Approval

All experiments were performed under IACUC protocol 18013: PI Vidhya Munnamalai, in compliance with the U.S. Department of Health and Human Services and reviewed by The Jackson Laboratory Institutional Animal Care and Use Committee.

## 7. Data Availability

The datasets used and/or analyzed during the current study are available from the corresponding author on request.

